# Distinguishing gene flow between malaria parasite populations

**DOI:** 10.1101/2021.01.08.425858

**Authors:** Tyler S. Brown, Aimee R. Taylor, Olufunmilayo Arogbokun, Caroline O. Buckee, Hsiao-Han Chang

**Affiliations:** Center for Communicable Disease Dynamics, Harvard T.H. Chan School of Public Health; Infectious Diseases Division, Massachusetts General Hospital; Infectious Disease Epidemiology and Ecology Lab, University of North Carolina School of Medicine; Institute of Bioinformatics and Structural Biology, National Tsing Hua University

## Abstract

Measuring gene flow between malaria parasite populations in different geographic locations can provide strategic information for malaria control interventions. Multiple important questions pertaining to the design of such studies remain unanswered, limiting efforts to operationalize genomic surveillance tools for routine public health use. This report evaluates numerically the ability to distinguish different levels of gene flow between malaria populations, using different amounts of real and simulated data, where data are simulated using parameters that approximate different epidemiological conditions. Specifically, using *Plasmodium falciparum* whole genome sequence data and sequence data simulated for a metapopulation with different migration rates and effective population sizes, we compare two estimators of gene flow, explore the number of genetic markers and number of individuals required to reliably rank highly connected locations, and describe how these thresholds change given different effective population sizes and migration rates. Our results have implications for the design and implementation of malaria genomic surveillance efforts.

## 1 Introduction

Measuring the extent to which malaria parasite populations are linked across different geographic locations (“connectivity”) can provide important guidance for the design and implementation of malaria control strategies. For example, estimates of connectivity can help predict the spatial dispersal of genetic variants associated with drug resistance. Patterns of connectivity are expected to differ with spatiotemporal variations in the population structure of parasites, mosquitoes, and humans, and thus differ across different epidemiological contexts.

Multiple approaches are available for measuring connectivity between malaria populations. Genomic surveillance approaches aim to directly measure gene flow between malaria parasite populations. Other measurements of connectivity, including estimates of human mobility [1], can provide indirect evidence of connectivity between malaria parasite populations, but such data are often limited in rural and outlying areas that may serve as reservoirs for continued malaria transmission [2].

The fixation index (*F_ST_*), a measure of population differentiation based on allele frequency variation, is widely used to assess gene flow between malaria parasite populations. Although methods for translating different amounts of differentiation to migration exist [3], a common approach used in malaria genomic surveillance studies is to consider malaria populations as somewhat differentiated if the confidence interval around their *F_ST_* estimate does not include zero [4, 5, 6, 7]. This binary approach is limited in situations where ranking weakly versus strongly connected populations is important.

There is currently no established guidance for the task of ranking different levels of connectivity between malaria populations, and multiple questions related to study design remain open. These include questions about different measures of gene flow and the data required to estimate them in different settings. Recent work suggests that measures based on identity by descent (IBD) may be superior to *F_ST_* for recovering connectivity between local populations [8]. Unlike *F_ST_*, IBD-based measures capture variation in relatedness introduced by recombination, and are thus likely better suited to measuring gene flow where variation caused by recombination exceeds that caused by drift. The properties of various estimators of *F_ST_* have been studied extensively [9]. Similarly, there is a rich and longstanding literature on IBD [10], including a recent study of the data requirements for estimating relatedness between malaria parasites [11]. Less is known, however, about how population summaries of inter-individual IBD estimates compare to *F_ST_* estimates when ranking different levels of connectivity between malaria parasite populations, or about how many individuals need to be sampled and how many genetic markers need to be genotyped per individual to reliably rank these estimates of gene flow in different settings.

Using real and simulated *P. falciparum* sequence data, we examine how the ability to rank estimates of *F_ST_* and an inter-population summary of IBD-based relatedness varies with the number of genetic markers (here we use data on single nucleotide polymorphims, SNPs) and the number of monoclonal *P. falciparum* samples. Using simulated data, we also examine how the ability to rank estimates of gene flow varies given different effective population sizes and rates of migration.

## 2 Methods

### 2.1 *P. falciparum* genomic data

We obtained *P. falciparum* whole genome variant call data, generated using the Genome Analysis Toolkit (GATK, [12]), from the MalariaGen database (Pf3k data release 5.1 [13]). We restricted our analysis to samples from the Greater Mekong Subregion (GMS), excluding more widely divergent *P. falciparum* populations from Sub-Saharan Africa and Bangladesh that are less relevant to the context of our analysis (i.e. genomic surveillance at the regional and national level), and including only individual sequences from clinical cases or survey participants. After excluding “un-callable” regions of the *P. falciparum* genome (i.e. highly repetitive or highly variable regions in which short-read based genotyping is unreliable [14]) and sites failing any GATK quality filter, we removed suspected polyclonal sequences and those with poor quality sequence data based on the proportions of heterozygous and missing SNP calls across all sites for each sample, and removed low-quality SNPs based on site-wise missingness and heterozygosity (similar to [15]). We first removed SNP sites for which > 1% of sequences had missing or heterozygous calls and then removed sequences for which > 5% SNP sites were missing or heterozygous. In addition, we removed two locations in Thailand (Sisakhet and Ranong) that each had less than 10 individual sequences available after these filtering steps. This filtering protocol yielded 472 individual sequences and 143,480 SNPs. Of the 143,480 SNPs, 418 had minor allele frequency estimates (computed using all 472 sequences pooled) > 0.35, where 0.35 is equal to a threshold used to select markers included in a widely-used malaria SNP barcode [16]. We restricted our analysis to these 418 SNPs.

Given evidence that a selective sweep on genes linked to artemisinin and piperaquine resistance (*kelch13* and *plasmepsin2-3*, respectively) substantially altered *P. falciparum* population structure across the GMS after 2012 [17, 18, 19], we included only sequences from 2009-2011 and excluded all sequences carrying the *kelch13* haplotype associated with this selective sweep (called KEL1 and identified using the haplotype scoring system in [19]). Lastly, we excluded two Pf3k study locations (Sisakhet and Ranong, Thailand) from the analysis that included < 10 *P. falciparum* individual sequences after removing post-2011 sequences and KEL1 mutants. Details on sample number and location are listed in Table 1. To evaluate whether estimated gene flow decays with distance between locations, we obtained estimated shortest road distances between Pf3k study locations from the Google Maps Distance Matrix API [20].

**Table 1:** County, location, and number of monoclonal *P. falciparum* sequences obtained from the Pf3k database [13], 2009-2011 with KEL1 mutants excluded as described in Methods.

| Country | Cambodia |  |  |  | Laos | Myanmar | Thailand | Vietnam |  |
| --- | --- | --- | --- | --- | --- | --- | --- | --- | --- |
| Location | Pailin | Pursat | Preah Vihear | Ratanakiri | Attapeu | Bago Division | Mae Sot | Bu Gia Map | Phuoc Long |
| Number of sequences | 10 | 104 | 60 | 126 | 59 | 12 | 37 | 42 | 22 |

### 2.2 Coalescent simulation of gene flow between malaria parasite populations

We used multi-population coalescent-based simulation [21] to extend our analysis to different settings, with specific focus on how differences in migration rates and effective population sizes (*N*_e_, which is known to correlate with transmission intensity of *P. falciparum* [22]) influence the ability to rank estimates of gene flow. We simulated sequence data for a metapopulation with 10 populations, with effective population sizes chosen within the range of values reported for locally-sampled malaria populations in low- and high-transmission settings (*N_e_* = 100 and 800, respectively) [22]. To account for facultative self-fertilization [23], an estimate of the recombination rate based on a cross between genetically distinct *P. falciparum* parasites, *ρ* = 1*E* − 7 recombination events per site per generation [14, 24], was scaled by *ϕ* = 0.044, where the value of *ϕ* was chosen by matching summaries of the simulated data to summaries of the Pf3k data. This results in an effective recombination rate of *ρ* × *ϕ* = 4.36*E* − 9 recombination events per site per generation input into the coalescent model. The mutation rate was set at 6.82E-9 mutations per site per generation. A 10-by-10 symmetric matrix of between-population baseline migration rates was generated as follows. First, for each row of a 10-by-10 matrix, we drew migration rates from a Dirichlet distribution parameterized by a random permutation of the vector (1, 2, 3, 4, 5, 6, 7, 8, 9,10). We then added the matrix transpose and divided by two, such that migration rates between locations are symmetric (i.e. equal in both directions). Migration rates were varied between simulations by multiplying the baseline migration rate matrix by a constant *M* = 0.1 or 0.5. For each of four parameter sets (*M*=0.1 and *N*_e_=100, *M*=0.1 and *N*_e_=800, *M*=0.5 and *N*_e_=100, and *M*=0.5 and *N*_e_=800), we obtained 100 independent replicate simulations. For each simulation replicate, we sampled 100 individuals from each population and discarded all simulated markers whose minor allele frequency estimates were ≤ 0.35, where estimates were computed using data from all 1000 individuals (100 individuals per population across 10 populations). For consistency with the terms used to describe the Pf3k data, we hereafter refer to populations in the coalescent simulation as “locations”.

### 2.3 Estimating gene flow between malaria parasite populations

Following [11], we define relatedness, *r*, between two individuals as the probability that, at any locus across the genome, the alleles for both individuals are IBD. In this study “individual” refers to either sequence data obtained from a single monoclonal *P. falciparum* infection in the Pf3k data or sequence data from a single individual in the simulations. We estimate relatedness between pairs of individuals using *hmmIBD* [25], taking the output variable `fract_sites_IBD` as an estimate of inter-individual relatedness. Following [11], let 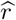 denote an inter-individual relatedness estimate. To summarize numerous relatedness estimates between individuals from different locations, let 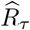 denote the proportion of between-location estimates greater than some specified threshold, *τ*, where 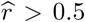 is considered highly related, and 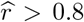 is considered nearly clonal. We estimate inter-location differentiation using the Weir and Cockerham estimator of *F_ST_* [26]. Let 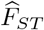 denote an estimate of inter-location differentiation. We refer to 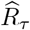 and 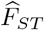 as estimates of gene flow since we expect gene flow between malaria parasite populations to correlate with both (positively with 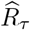 and negatively with 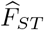). Other drivers of genetic variation (e.g. selection) can contribute to both.

We calculate 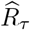 and 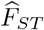 using different numbers of individuals per location, *n*, and different numbers of SNPs, *p*, and benchmark against estimates calculated using all individuals per location, *N*, and all available SNPs, *P*. Let 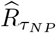 and 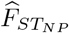 denote the estimates calculated using all available data. For the Pf3k data, 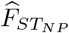 and 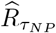 are calculated using all 418 SNPs with minor allele frequency estimates > 0.35 and, for each location pair, the maximum number of individuals available for each location (e.g., 104 individuals from Pursat, Cambodia, and 37 individuals from Mae Sot, Thailand). For the simulated data, we calculate these values using all 100 sampled individuals from each location and all available SNPs with minor allele frequency estimates > 0.35 (the number of which varies between simulation replicates). We stratify comparisons between location pairs using the differences between gene flow estimates calculated using all available data per location pair; let 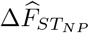 and 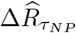 denote these differences.

### 2.4 Estimating the ability to rank different levels of gene flow

We next consider the ability to rank estimates of gene flow between location pairs computed using different numbers of individuals per location, *n*, and SNPs, p. Let 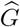 denote an estimate of gene flow, either 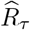 or 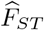. For a pair of location pairs, e.g. the kth location pair and the *k* + 1th location pair, 100 estimates of gene flow per location pair are generated as follows. First, draw uniformly at random without replacement a set of *p* SNPs (without replacement because SNP positions feature in the estimation of r); then for *i* =1, 2,.., 100, draw uniformly at random with replacement *n* individuals per location, and then compute 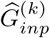 and 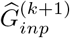 where the superscripts denote the location pairs and the subscripts indicate that the estimates were computed using data on the ith randomly-sampled set *n* individuals per location and the set of *p* SNPs. Given all 100 estimates of gene flow per location pairs *k* and *k* +1, the goal is to compute the proportion of pairs of estimates that are ranked correctly. An approximation of the proportion ranked correctly, PRC_*np*_, is computed as follows,

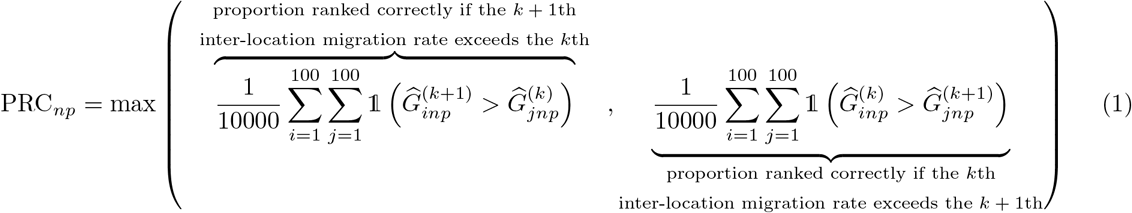

where 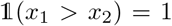 if *x*_1_ > *x*_2_ and 0 otherwise. PRC_*np*_ is an approximation of the true proportion ranked correctly because, instead of comparing to the actual migration rates, which are unknown for the Pf3K data, we take the maximum proportion ranked correctly given the two possibilities: either the *k* + 1th inter-location migration rate exceeds the *k*th or vice versa; see equation (1). PRC_*np*_ ∈ [0,1] with PRC_*np*_ = 1 when the ranges of gene flow estimates per location pair do not overlap and 0 when all the gene flow estimates are equal (i.e. when 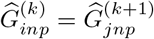 for all *i,j* = 1,…, 100).

Our analysis focuses on the specific task of ranking location pairs with the highest shared gene flow versus those with lower shared gene flow, which in practice would allow for stratification of locations into highly connected and less connected groups. We chose to designate highly connected location pairs as those with the five largest 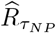 values when 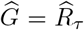, and five smallest 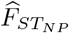 values when 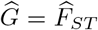. The remaining location pairs (31 in the Pf3k data and 40 in simulated data) are consider less connected. For the Pf3k data, for each value of *n* and *p*, we generate 155 PRC_*np*_ values, one for each comparison between 5 highly connected location pairs and 31 less connected pairs. As such, plots of PRC_*np*_ based on real data feature 155 points per *n* and *p* combination. These are are coloured by 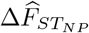 and 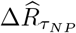. For the simulated data, we consider 200 comparisons between 5 highly connected and 40 less connected location pairs.

Using simulated data, for a given combination of *n* and *p*, we generate 100 values of PRC_*np*_ per pair of location pairs: one for each of the 100 independently run replicate simulations, except where the number of SNPs with estimated minor allele frequency > 0.35 is less than *p*, in which case we compute an additional PRC_*np*_ using a randomly selected replicate whose SNP count is adequate. As such, plots of PRC_*np*_ for highly versus less highly connected population pairs based on simulated data feature error bars based on 20000 values of PRC_*np*_. In an additional set of analyses using the simulated data, we consider the proportion ranked correctly for all 990 pairs of location pairs (ten choose two choose two). For these analyses, we calculate 99000 values of PRC_*np*_ for each value of *n* = 5, 10,…, 95, 100; PRC_*np*_ values are parsed into bins by *n* and 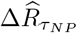 (or 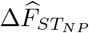) and the mean of the values of PRC_*np*_ for each bin are displayed on a contour plot.

### 2.5 Code availability

Code for generating simulated data and calculating PRC_*np*_ values is available at https://github.com/tsbrown-git/malariarankedgeneflow

## 3 Results

### 3.1 Relatedness and differentiation in real and simulated *P. falciparum* sequence data

Figure 1 shows the distribution of between-location values of 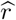 by study location in the Pf3k data. Nearly clonal between population pairs likely result from recent migration events (where the time since migration is short enough that there is limited or no out-crossing between imported individuals and the receiving population) and as such provide an important indicator of ongoing, epidemiologically-relevant connectivity between *P. falciparum* populations. Between population sample pairs with lower 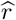 values (for example 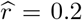) reflect more distant gene flow between malaria populations and thus may not capture migration events most relevant for quantifying recent connectivity between populations.

**Figure 1:**
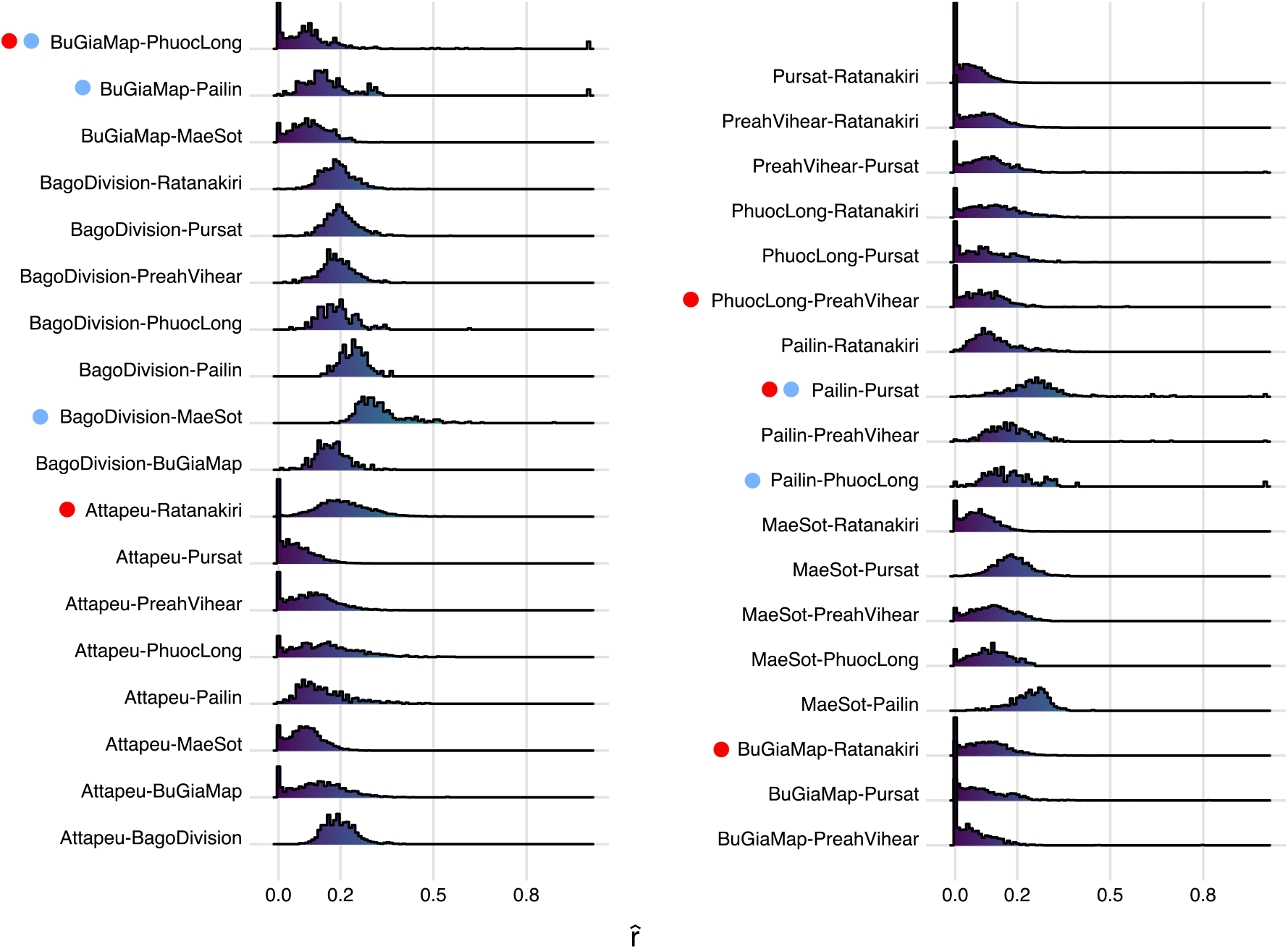
Density plots of 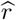 for between-location pairs of individuals at nine study locations from the Pf3k data. For each of the nine choose two (36) location pairs, density is shown on the vertical axis. Location pairs with the five lowest pairwise 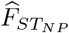 values are annotated with red circles and those with the five highest 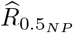 are annotated with blue circles.

Differences between distributions of 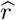 values likely reflect differences in sampling procedures and *P. falciparum* population structure between Pf3k study locations. Indeed, differences are evident in the within-location distributions for 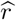 by study site (Supplemental Figure S1 A), where we observe substantial heterogeneity in the proportion of clonal pairs 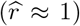 in different study locations, which could reflect both true differences in levels of within-population diversity and/or sampling bias (for example, if samples were collected from individuals infected with the same clone during an outbreak or other shared transmission event).

Supplemental Figures S1 B and S2 show the distributions of within-population 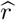 and between-population 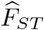 values computed using simulated data. Simulations with M=0.5 and *N*_e_ = 100 provide the best approximation of within-population 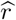 for the Pf3K data set (top row, supplemental Figure S1 A) and simulations with *M* = 0.1 and *N*_e_ = 800 most closely match the distribution of between-population 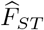 values observed in the Pf3k values (top rows, Supplemental Figure S2).

### 3.2 Highly connected location pairs

The five most highly connected location pairs in the Pf3k data, as measured by their 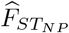 and 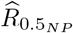 values, are shown in Figure 2 and annotated on Figure 1. Three location pairs (Bu Gia Map – Pailin, Bago Division – Mae Sot, and Pailin – Phuoc Long) with high values for both 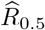 (Figure 2 B) and 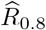 (Supplemental Figure S3) are not included in the set of five location pairs with the lowest 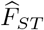 values (Figure 2 A), suggesting that 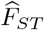 may fail to identify some location pairs with higher shared gene flow attributable to recent migration events.

**Figure 2:**
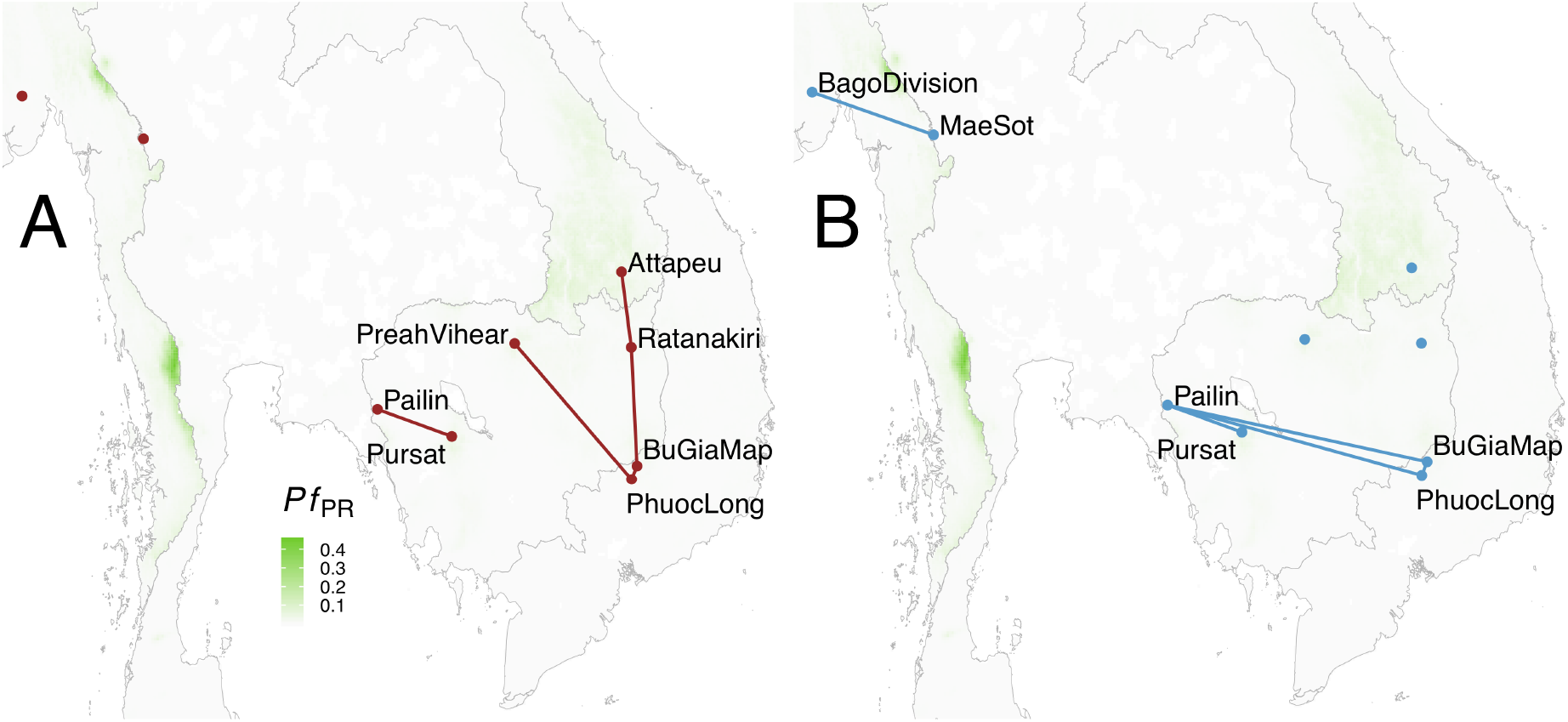
Pf3K location pairs with lowest between-population differentiation, 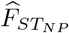 (A) and highest between-population relatedness, *R*_0.5,*NP*_ (B) measured using *n* = 418 SNPs and the maximum number of individuals in each population. Background map is colored by the estimated *P. falciparum* parasite rate, *Pf_PR_*, for 2012. [31]

**Figure 3:**
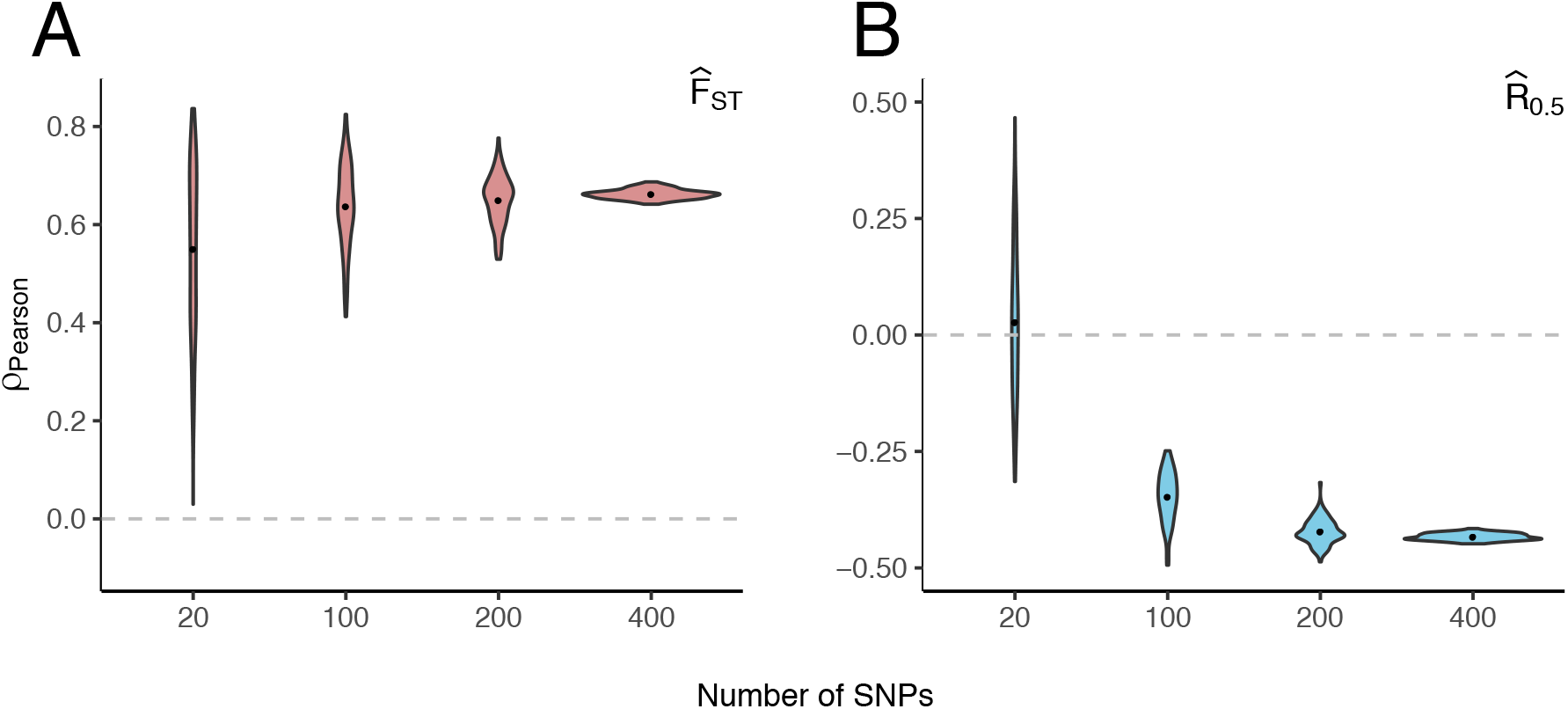
Pearson’s correlation coefficient *ρ* for either (A) 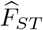 (B) or 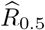 versus shortest road distance between Pf3K locations by number of SNPs. Violin plots represent distribution of Pearson’s *ρ* values for 100 independently sampled SNP sets, each of size *p* = 20, 100, 200, or 400.

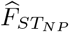 values are positively correlated with shortest road distance between study locations, i.e. genetic differentiation increases with distance between locations; 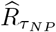 values are negatively correlated with road distance given *τ*=0.5 (Figure 2) and 0.8 (Supplemental Figure S3), consistent with decreasing between-population proportions of highly related and nearly clonal parasites with increasing distance. No correlation with distance is observed for *τ*=0.2 (Supplemental Figure S3).

### 3.3 The ability to distinguish location pairs

#### 3.3.1 The ability to rank highly connected Pf3k location pairs

We first evaluated how well gene flow estimates for location pairs with the highest levels of gene flow (based on 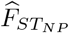 and 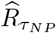) can be ranked versus those with comparatively lower gene flow. Figure 4 shows the ability to rank highly versus less highly connected location pairs in the Pf3k data by number of SNPs and number of individuals. In general, PRC_*np*_ is lower for comparisons between location pairs with smaller values of 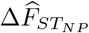 and 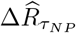. For both 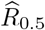 and 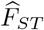, the ability to identify highly connected population pairs (Figure 4) is limited for small numbers of individuals (*n* = 10 and 12) even with larger numbers of SNPs. With *n* = 22 and *p* = 200 and 400, 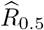 identifies highly connected population pairs with PRC_*np*_ > 0.95 in the majority of comparisons between population pairs (85% and 95%). 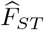 identifies highly connected population pairs less reliably when the same number of SNPs (*p* = 200 and 400) and individuals (*n* = 22) are used (70% and 75%). The ability to identify highly connected population pairs is decreased when using a more stringent threshold for relatedness, *τ* = 0.8: only 79.2%, of comparisons between highly connected and less highly connected pairs can be ranked correctly with PRC_*np*_ > 0.95 with *n* =22 and *p* = 200 and 400 SNPs (Figure S4). However, 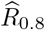 outperforms 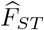 for *n* = 22 and *p* = 24,100 and 200.

**Figure 4:**
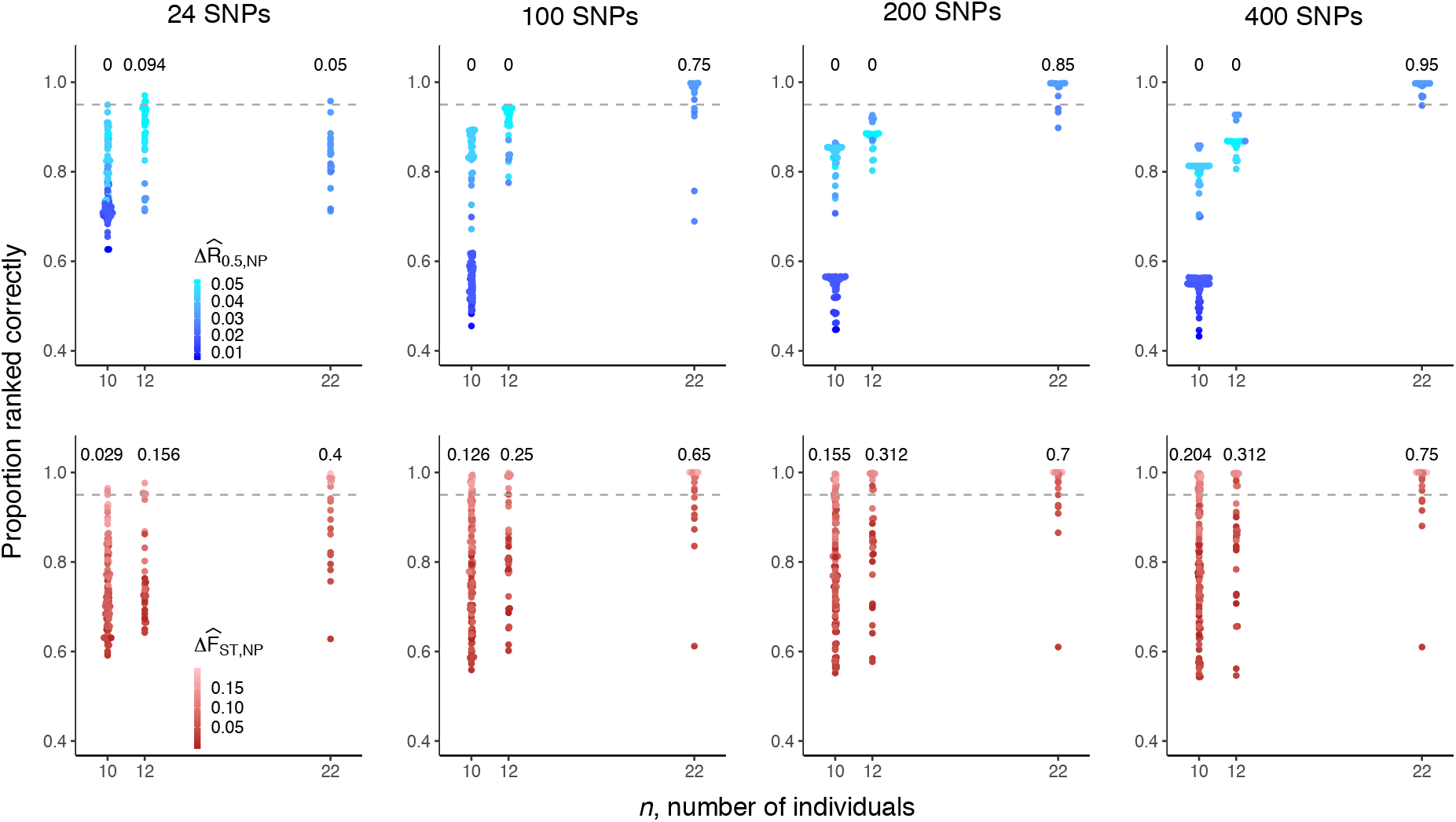
The ability to rank highly versus less highly connected population pairs by number of SNPs and number of individuals using 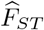 or 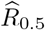. The proportion ranked correctly (PRC_*np*_) is plotted against the maximum number of individuals available for each comparison between population pairs. Points are colored by 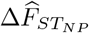 or 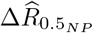, the difference between gene estimates computed using all available data. For each value of *n* and *p*, 155 total PRC_*np*_ values are plotted (i.e. the number comparisons between 5 location pairs with highest values for 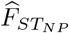 or 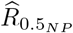 and the remaining 31 location pairs with lower gene flow). The horizontal dashed grey line demarcates comparisons between location pairs where PRC_*np*_ is less than or greater than 0.95; the proportion of location pairs with PRC_*np*_ > 0.95 is annotated above each number of individuals.

#### 3.3.2 The ability to identify highly connected simulated location pairs

We observe similar requirements for number of SNPs and number of individuals using sequence data simulated under different effective population sizes and migration rates. Overall, the ability to rank highly connected versus less highly connected location pairs is lower for larger migration rates and effective population sizes (Figures 5 and 6). Similar to observations from the Pf3k data set, the ability to rank is strongly influenced by the number of individuals sampled, *n*, and low ability to rank with small *n* is not substantially improved by using larger numbers of markers, p. Across simulations, the proportion of pairs of gene flow estimates that are ranked correctly is most limited for *n* < 40 individuals. For values of *n* comparable to those examined in the Pf3k data (i.e., 5 < *n* < 30), we find close correspondence between PRC_*np*_ values in simulations with *N*_e_=100 (Supplementary Figure S5). In the simulations with *N*_e_=800 and M=0.5 (i.e., the largest population size and migration rate), neither 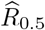 nor 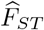 achieve PRC_*np*_ > 0.95, even with the maximum number of *p*=200 SNPs and n=100 individuals (right columns, Figures 5 and 6). This observation suggests that in some settings, with large effective populations sizes and between-population migration, the reliable distinction of gene flow between *P. falciparum* populations is not possible using either 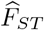 or 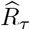.

**Figure 5:**
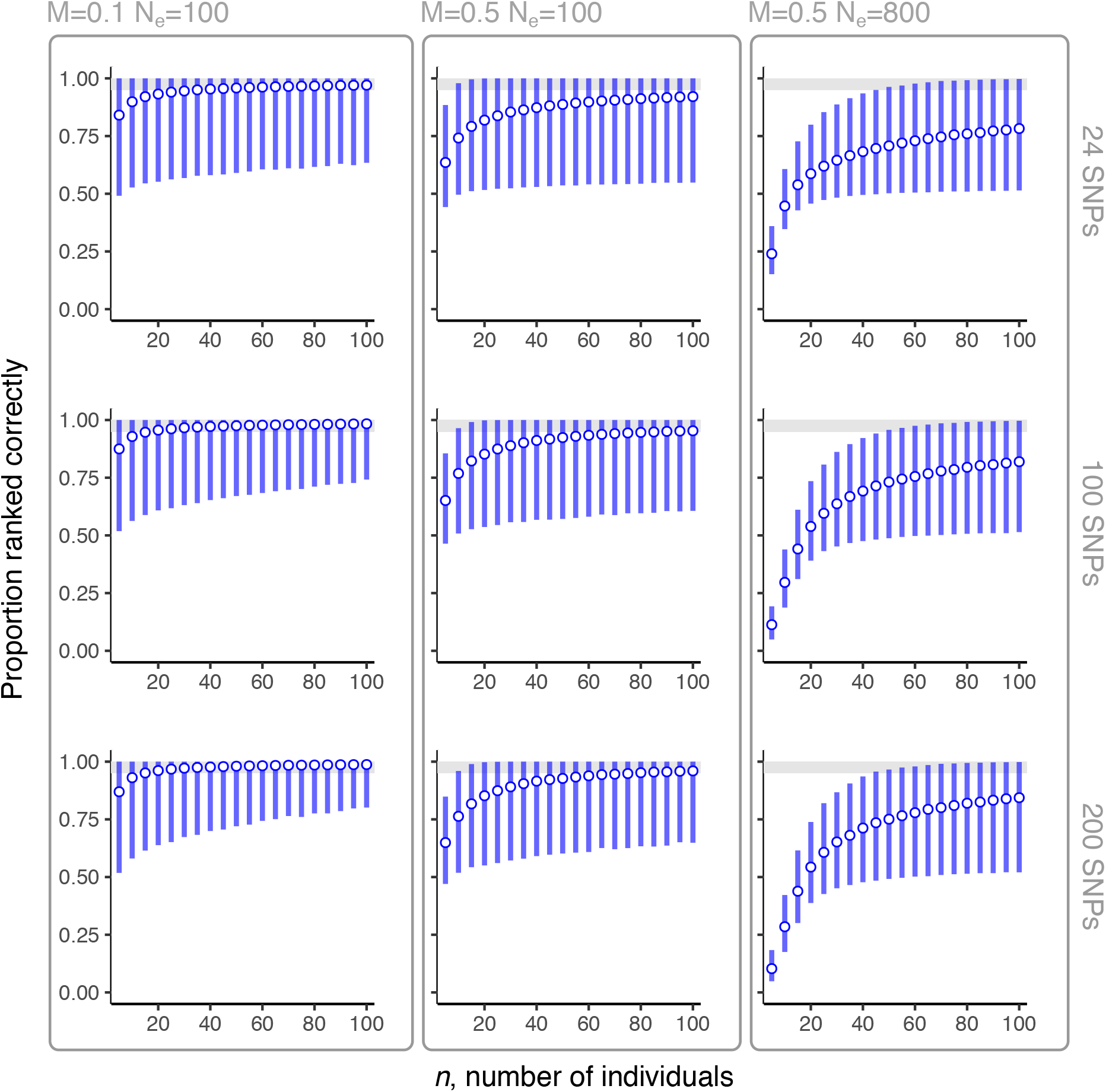
The ability to rank highly versus less highly connected population pairs by number of SNPs and number of individuals using 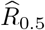 for coalescent-simulated sequence data. Results for 24, 100, 200 SNPs, tested over low and high migration scenarios (M=0.1 and M=0.5, respectively) and small and large effective population sizes (*N*_e_=100 and *N*_e_=800). Points show the mean of 20000 PRC_*np*_ estimates with error bars showing the 0.025 and 0.975 percentile values. Grey area demarcates where PRC_*np*_ > 0.95.

**Figure 6:**
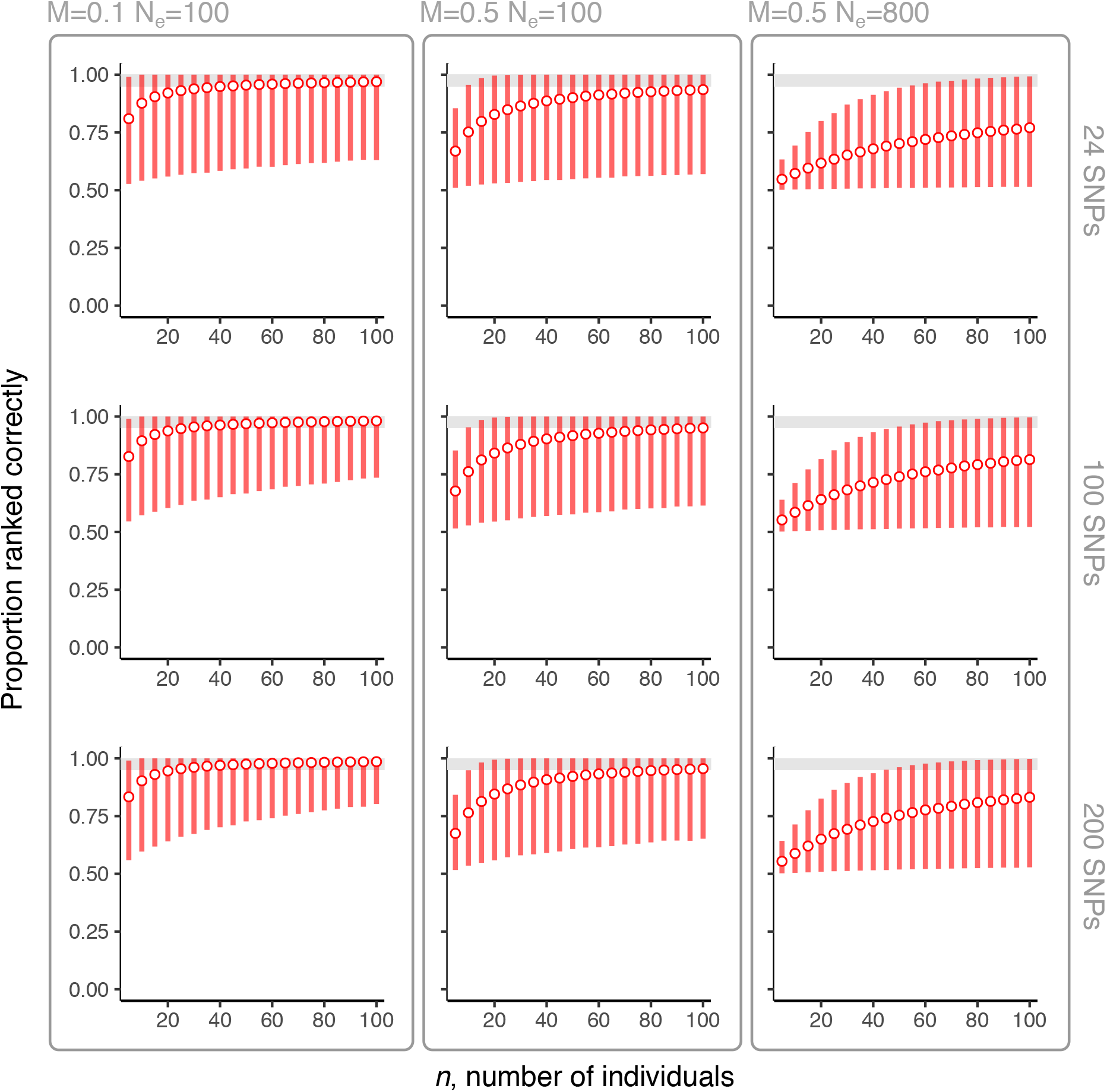
The ability to distinguish highly versus less highly connected population pairs by number of SNPs and number of individuals using 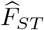 for coalescent-simulated sequence data. Results for 24, 100, 200 SNPs, tested over low and high migration scenarios (M=0.1 and M=0.5, respectively) and small and large effective population sizes (*N*_e_=100 and *N*_e_=800). Points show the mean of 20000 PRC_*np*_ estimates with error bars showing the 0.025 and 0.975 percentile values. Grey area demarcates where PRC_*np*_ > 0.95.

#### 3.3.3 The ability to rank simulated location pairs

To understand more generally how the proportion of pairs of gene flow estimates are ranked correctly varies as a function of *n, p*, and 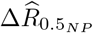 or 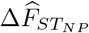, we calculated PRC_*np*_ for all 990 possible comparisons between location pairs using data simulated under different effective population sizes and migration rates (Figures S6 and S7). In simulations with smaller population sizes (*N*_e_ = 100), which approximate the distributions of within-location 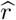 values but not between-location 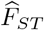 values observed for the Pf3k data (Figures S1 and S2), PRC_*np*_ >95% over a wide range of *n* values (for both 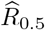 and 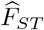), and this range of *n* values with PRC_*np*_ >95% increases with increasing p. For *p* = 100 and *p* = 200, PRC_*np*_ < 0.95 when a small number of individuals are compared (*n* < 30) or for comparisons between location pairs with small values for 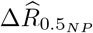 or 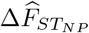. For *p* = 24 SNPs, larger numbers of individuals are required for PRC_*np*_ > 0.95 and only location pairs with relatively larger 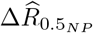 and 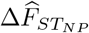 values have PRC_*np*_ > 0.95. For *N*_e_=800 or *M*=0.5, we observed a smaller number of comparisons between pairs with PRC_*np*_ >95% for PRC_*np*_ calculated using either 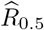 (Figure S6) or 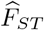 (Figure S7). With *N*_e_=800, *M*=0.5, and *p*=24, there are no comparisons between location pairs with PRC_*np*_ >95% when PRC_*np*_ is calculated using 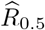, and a very small proportion of comparisons between locations pairs with PRC_*np*_ >95% when PRC_*np*_ is calculated using 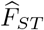.

## 4 Discussion

Genomic and genotype-based methods have become increasingly important tools for malaria surveillance and control, providing valuable information on the impact of disease control interventions [22], the geographic dispersal of antimalarial drug resistance [27], and connectivity between regional and local parasite populations [8], among other critical insights. There is an ongoing effort to operationalize these tools for use in routine disease surveillance activities, and developing the evidence base around the appropriate use of these tools is a central scientific and public health priority. In this study, we used genome sequence data from naturally-occurring *P. falciparum* infections and simulated sequence data using parameters associated with different epidemiological conditions to examine a specific but potentially important use case in malaria genomic surveillance, i.e. ranking gene flow between different *P. falciparum* population pairs.

We highlight four main observations from our analysis: (1) Highly related and nearly clonal between-location sample pairs provide an important marker for recent parasite migration between populations. Given a sufficient number of SNPs (*p* > 100) and individual samples (*n* ≥ 22), 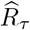 with *τ* = 0.5 and 0.8 captures decay with relatedness over distance; 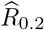 fails to capture decay in relatedness over distance, likely because the lower threshold obscures the signal from more highly related sample pairs and thus does not capture recent migration events. (2) Consistent with prior studies [8, 11], the ability to rank gene flow is lower with 24-SNP datasets compared to 100- or 200-SNP datasets. Performance is substantially lower when using 24-SNP versus 100-SNP datasets, but only marginally higher when using 200-SNP versus 100-SNP datasets; (3) The number of individuals sampled from each location (*n*) is an important determinant of the ability to rank gene flow. In both the *P. falciparum* sequence data and in simulations with higher and lower specified migration rates and smaller and larger population sizes, we observe markedly decreased proportions of pairs of estimates that are ranked correctly when *n* is less than 40 individuals; (4) High levels of migration and large effective population sizes may preclude reliable ranking of gene flow between populations, even when using larger numbers of SNPs or sampling large numbers of individuals.

There are multiple limitations that are important for understanding these findings in context. First, the *P. falciparum* sequence data we obtained from the Pf3k database includes only a small number of individuals from certain locations (for example, Pailin, Cambodia and Bago Division, Myanmar), such that we are able to evaluate the proportion of pairs of gene flow estimates that are ranked correctly using real *P. falciparum* sequence data over only a limited, discrete set of values for *n*. Second, our analysis of real *P. falciparum* sequence data is limited to samples collected over a two-year period (2009-2011) in a relatively small number of locations in Southeast Asia, and observations from this dataset are not generalizable beyond the epidemiological and ecological conditions specific to this context. Third, our analysis is likely influenced by biases inherent to sample collection in each Pf3k study location, making it difficult to disambiguate whether observed patterns in relatedness and differentiation are due to underlying *P. falciparum* population structure or sampling artifact (for example, if samples were collected from clinical patients infected with the same clone during an outbreak or other shared transmission event).

We sought to address some of these limitations via coalescent simulation, allowing for numerical evaluation of PRC_*np*_ over a wider range of parameters, specifically those intended to approximate the genetic characteristics of larger *P. falciparum* populations with higher levels of inter-population migration. Use of coalescent-simulated sequence data inherently avoids the biases in sampling from naturally-occurring *P. falciparum* populations described above and allows for the ground truth regarding migration rates and population sizes to be known (i.e. they are pre-specified as simulation parameters). Parameters used for coalescent simulation are subject to potential mis-specification, particularly those parameters where there is uncertainty about their true values in naturally-occurring *P. falciparum* populations (for example, recombination rate). Although we do not statistically infer model parameters, we note that, for some parameter combinations, the coalescent simulations used in our analysis generate data whose distributions of between-location 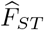 values and within-location 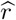 values are comparable to distributions computed using the Pf3k data.

There are other aspects of our approach that may limit generalizability to real *P. falciparum* populations in the field. For example, we have assumed the ability to sample individuals evenly across populations, and that sampling by default captures every population in the metapopulation; in real-life genomic surveillance applications, sampling is likely to be unbalanced and imperfect, such that individuals from certain populations may be undersampled or not captured at all. Lastly, coalescent simulation assumes perfect overlap of survey catchment area and population, whereas in real-life surveillance applications, it is more difficult to ascribe a known geolocation to each individual infection (given that individuals may move location between infection and medical diagnosis or survey participation) and thus the task of assigning individual infectious to specific locations involves much more uncertainty.

Lastly, we highlight four important technical issues where further work is needed. First, in our estimation of the proportion of pairs of gene flow estimates that are ranked correctly (equation (1)), we assume that the location pair that most often has a larger gene flow estimate is equal to the location pair that truly has greater gene flow. This assumption is unlikely to hold when differences in gene flow are small or the data are limited (given that uncertainty around inter-individual estimates of relatedness is large for small *p* [11]). For simulated data, this issue could be circumvented by consideration of the migration rates used to simulate the data. For Pf3k data, we could use gene flow estimates computed using all available data as best estimates of the ground truth. Second, when *τ* is high and *n* is small, the proportion ranked correctly for 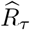 is low because many inter-population 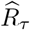 =0 for both location pairs. This issue may be circumvented by using threshold-free approaches (for example, using Wasserstein distances, as in [28]). Third, the SNP ascertainment scheme used in our analysis (including only SNPs with minor allele frequency estimates ≥ 0.35) could impact our ability to rank *F_ST_* estimates, e.g. by limiting their effective range if high minor allele frequencies enrich for markers under balancing selection. Meanwhile, for relatedness estimation, markers with high minor allele frequencies increase precision [11]. Fourth, the Weir and Cockerham estimator for *F_ST_* can perform poorly when the number of individuals is small [29], as they are here, or imbalanced [30]. An alternative estimator, e.g. the Hudson estimator, may perform more favourably [29, 30].

In conclusion, we have examined the numerical properties of 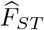 and 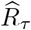 as estimates of population-level gene flow between *P. falciparum* populations, and outlined data requirements (specifically, number of individuals and number of genetic markers) for their use given simulation parameters that approximate different ecological and epidemiological conditions. Although this study is focused on a single genomic surveillance application (i.e. correctly ranking highly-connected location pairs over less highly connected ones), our work underscores an important limitation that applies across a wider range of surveillance applications, i.e. under certain epidemiological conditions (high migration between large populations) it likely becomes largely infeasible to reliably rank gene flow between *P. falciparum* populations, even with dense sampling of individuals and sequence data for large numbers of genetic markers. Additional scientific work is needed to understand how unbalanced sampling of individuals, incomplete sampling of metapopulations, and other exigencies of real-world genomic surveillance impact the ability to rank different levels of gene flow between *P. falciparum* populations in the field.

## 5 Supplemental Figures

**Figure S1:**
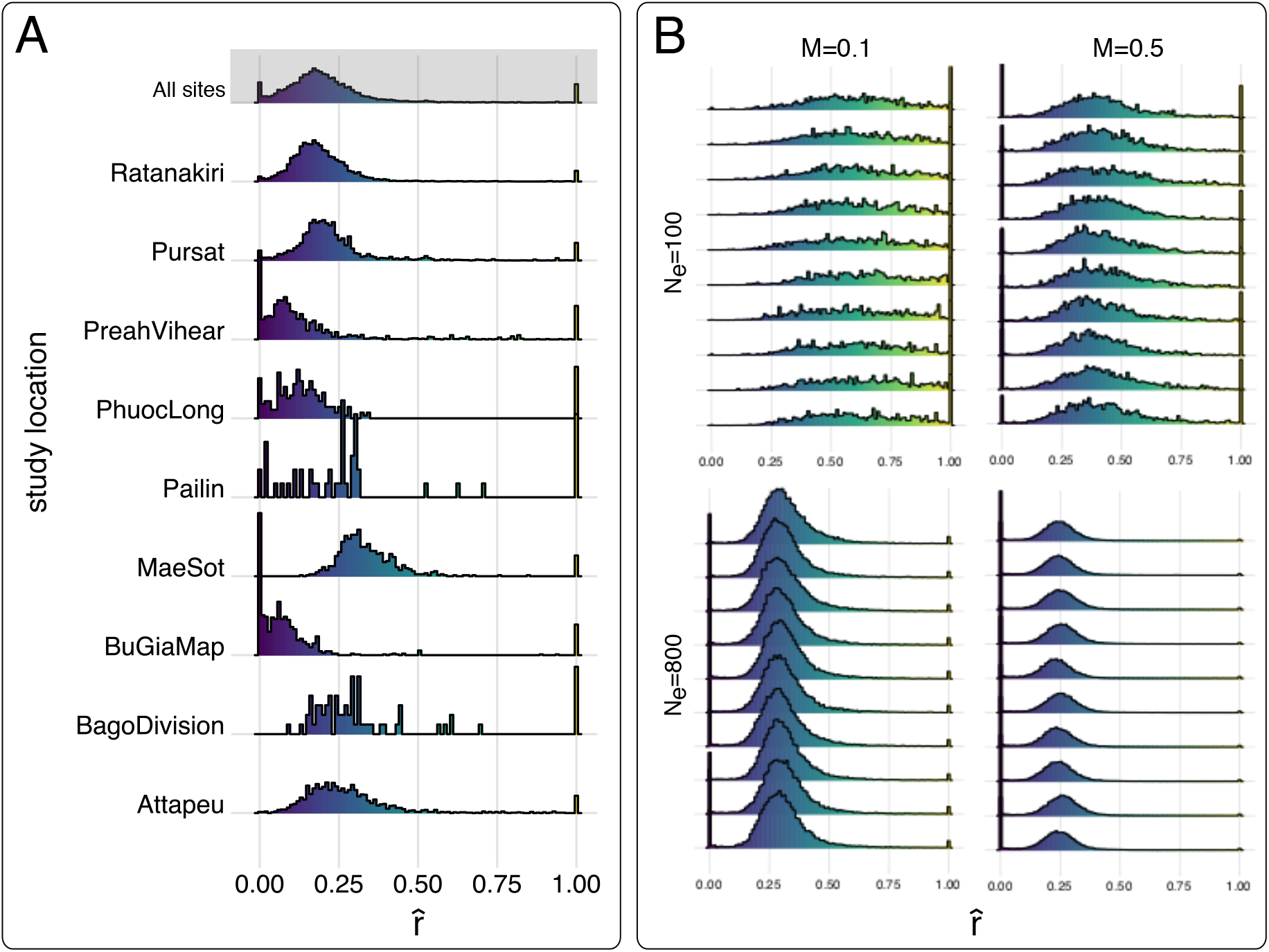
Density plots of within-population 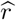 values for study location in the (A) Pf3k dataset and (B) coalescent simulation. Density is shown on the vertical axis. Each panel in B displays the distribution of all within-population 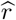 values (corresponding to the “all sites” distribution for the Pf3k dataset in Panel A) for ten representative simulation replicates given different levels of migration (M=0.1 or M=0.5) and effective population sizes (*N*_e_=100 or *N*_e_=800).

**Figure S2:**
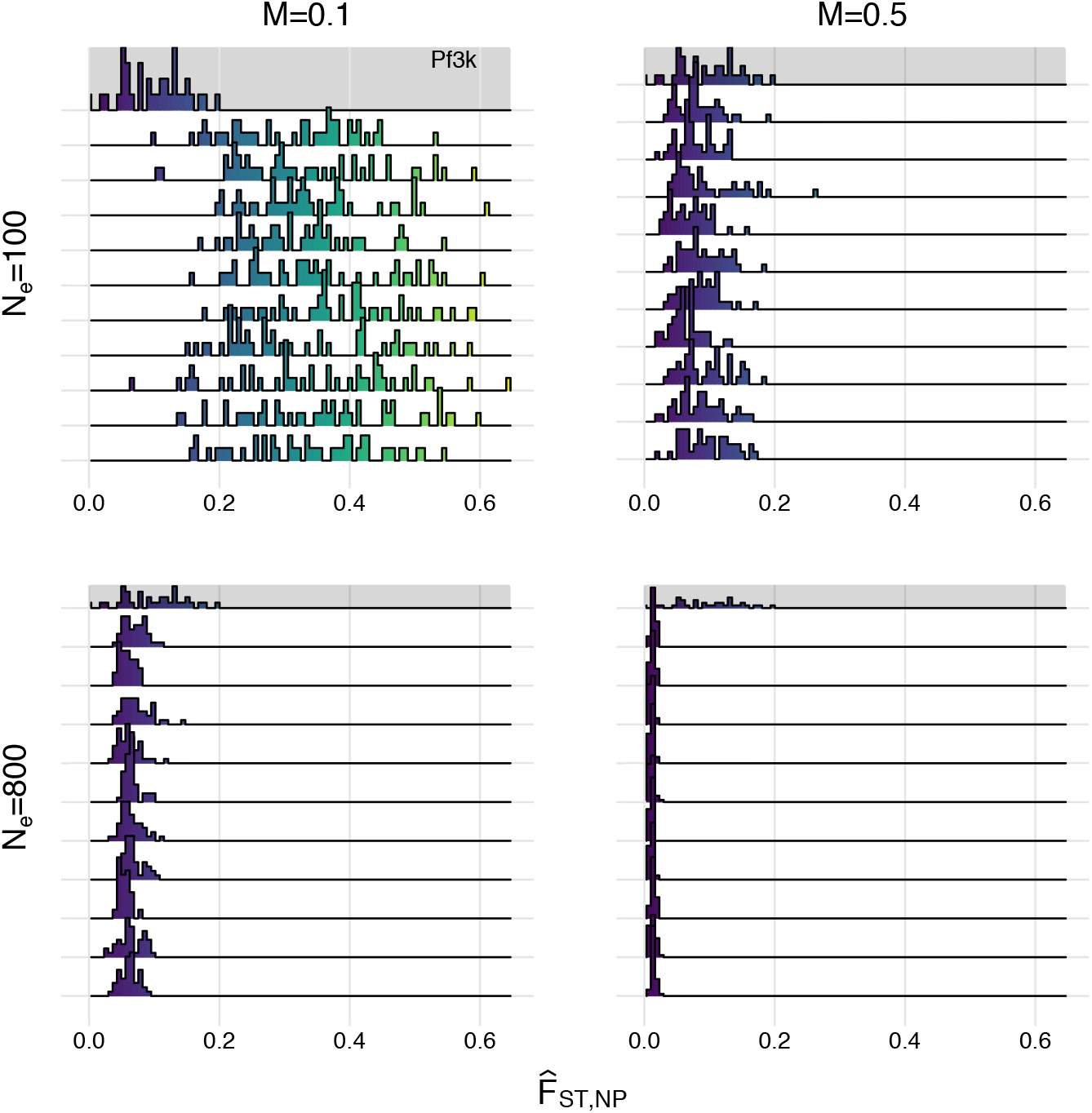
Density plots of 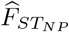 values from coalescent simulations. Panels each display ten representative simulation replicates under different levels of migration (*M*), and population sizes, *N*_e_. The distribution of pairwise estimates of 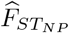 from the Pf3k data is highlighted in grey at the top of each panel. It is repeated four times because the range of density axes vary with M and *N*_e_.

**Figure S3:**
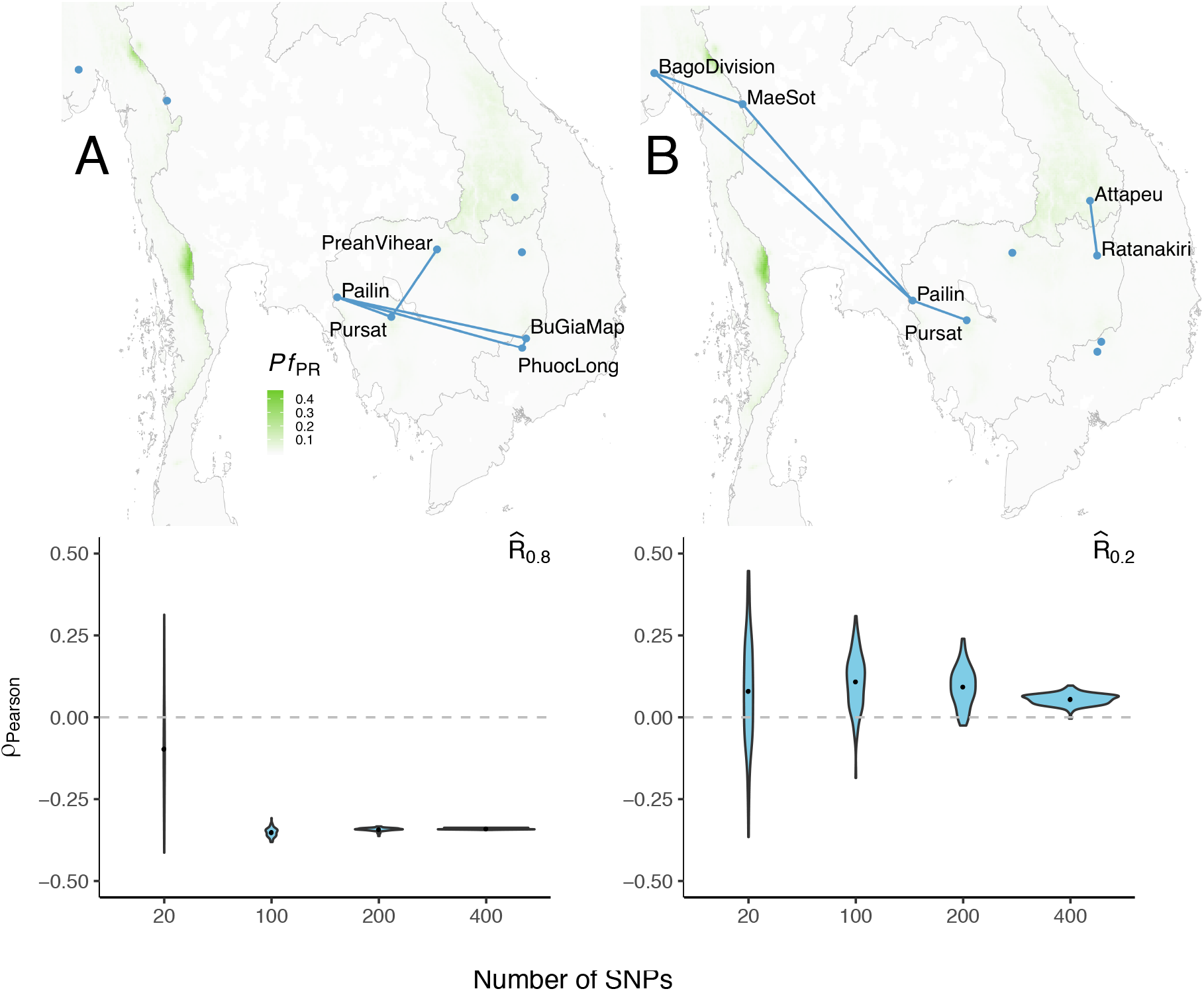
Top row: Location pairs with highest between-population relatedness, 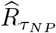 (measured using *p* = 418 SNPs and the maximum number of individuals in each population), for *τ* = 0.8 (A) and *τ* = 0.2 (B). Bottom row: Pearson’s correlation coefficient *ρ* for 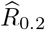 and 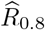 versus shortest road distance between locations by number of SNPs. Violin plots represent distribution of Pearson’s *ρ* values for 100 independently sampled SNP sets, each of size *p* = 20, 100, 200, or 400.

**Figure S4:**
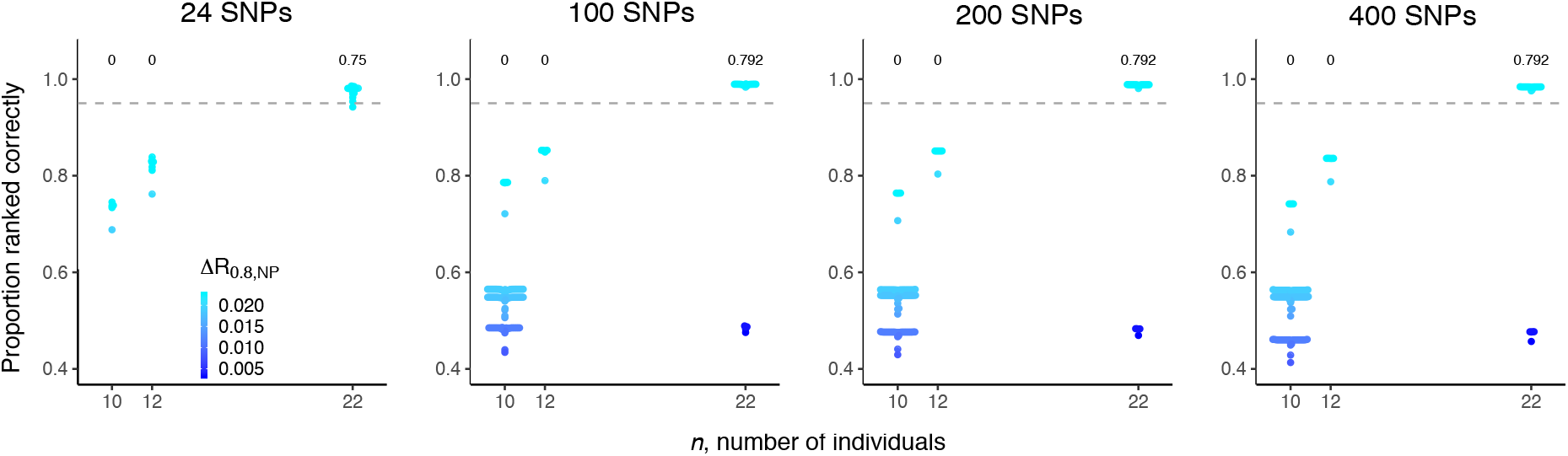
The ability to rank highly versus less highly connected population pairs by number of SNPs and number of individuals using 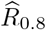. PRC_*np*_ is plotted against the maximum number of individuals available for each comparison between population pairs. Points are colored by 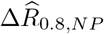. The grey line demarcates comparisons between location pairs with PRC_*np*_ < or > 0.95 and the proportion of all comparisons with the ability > 0.95 is annotated above each number of individuals.

**Figure S5:**
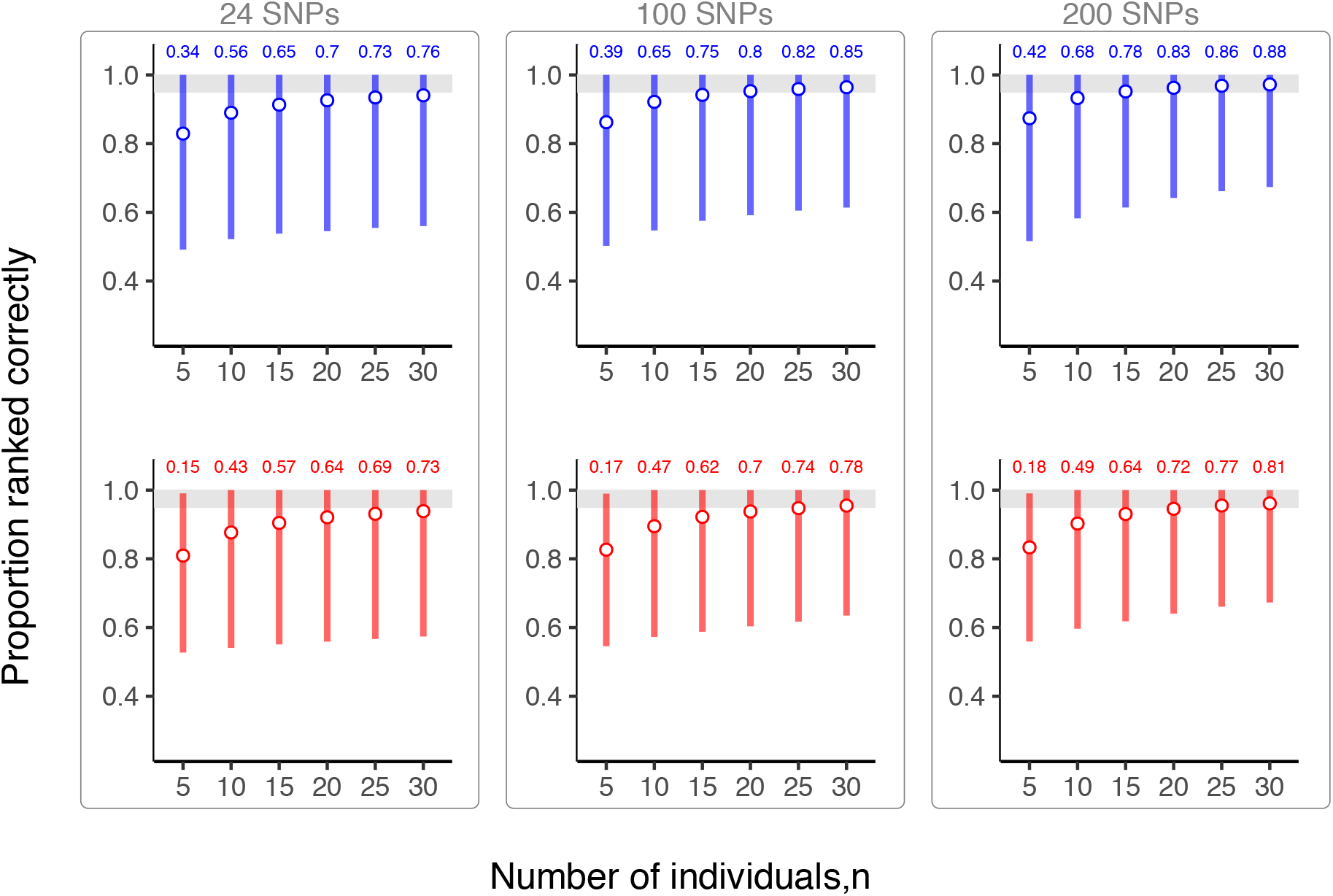
The ability to rank highly versus less highly connected population pairs for simulating sequence data using 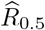 (upper row, blue) or 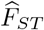 (bottom row, red). We show PRC_*np*_ values calculated using *n* = 5 to 30 sequences simulated with *N*_e_=100 and *M* = 0.1. Circles show mean proportion ranked correctly for all pairwise combinations between location pairs with highest connectivity (5 total) and all 40 other location pairs with lower connectivity across 100 simulation replicates. Error bars show the 0.025 and 0.975 percentile values for all 20000 PRC_*np*_ computed. The grey area demarcates where the region where PRC_*np*_ ≥ 0.95. For each *n*, proportions of PRC_*np*_*valuesgreaterthan0.95areannotatedatthetopoleachpanel*.

**Figure S6:**
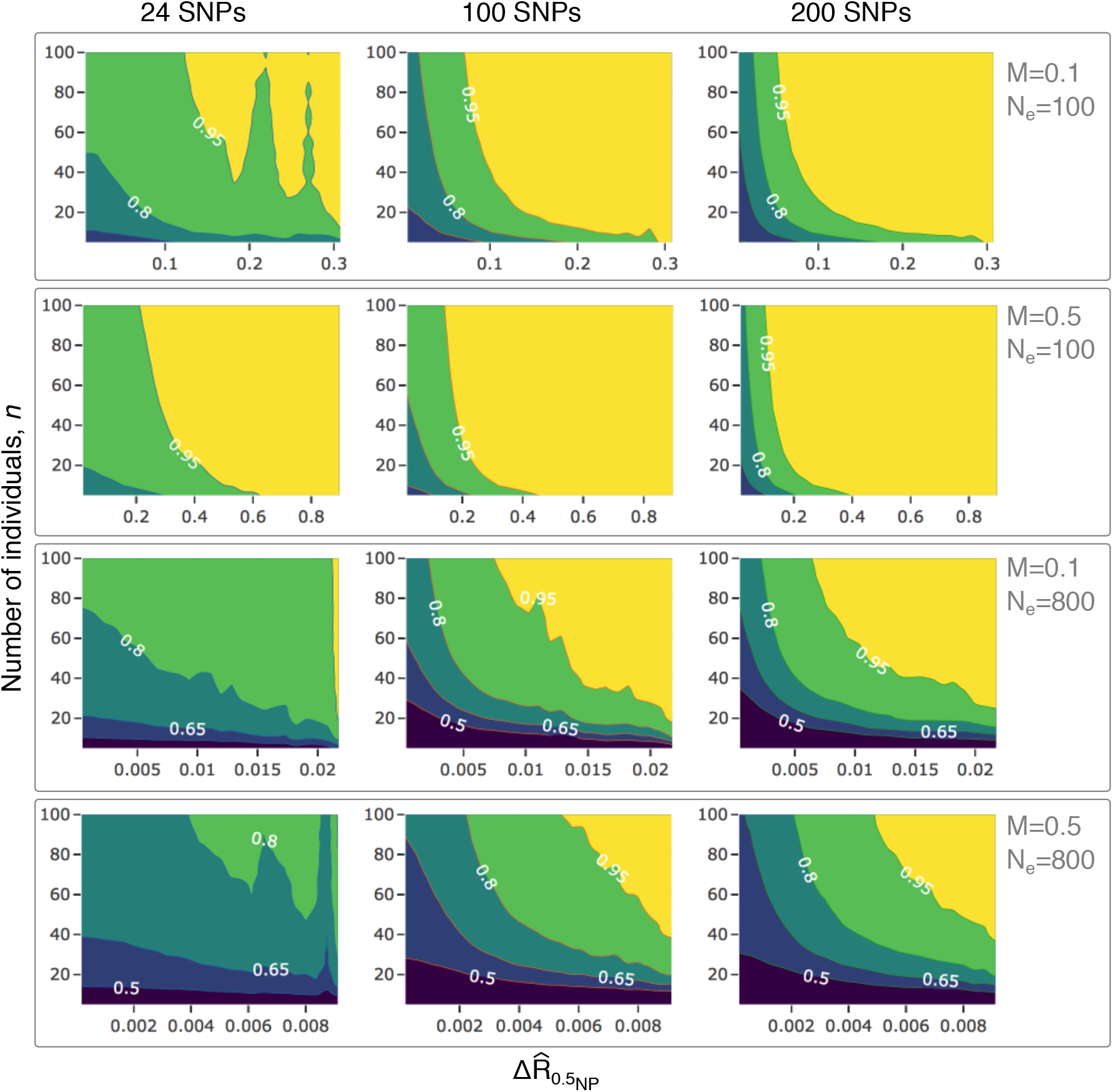
The ability to rank location pairs for coalescent-simulated data at different simulated population sizes and migration rates using 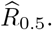 Contour plots show the ability to rank between location pairs by 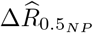 (horizontal axis), the absolute difference between 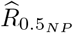 values for the two location pairs under comparison, and the *n*, the number of individuals sampled from each location (vertical axis). Each point represents the mean value for 100 independent values for PRC_*np*_, as described in the methods. *M* is a scaling factor for the distribution of pairwise migration rates between populations. *N_e_* is the effective population size specified for each coalescent simulation.

**Figure S7:**
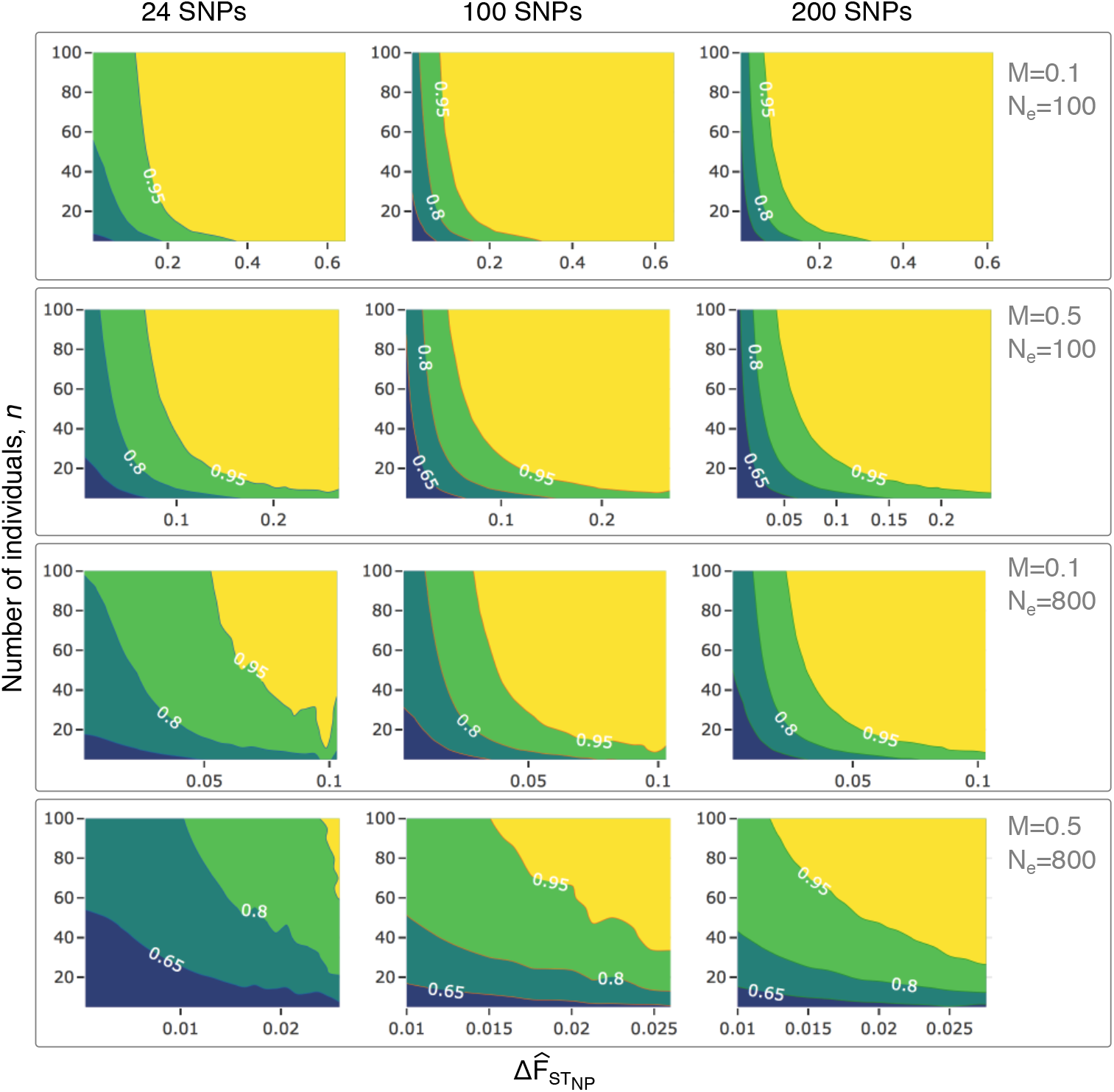
The ability to rank location pairs for coalescent-simulated data at different simulated population sizes and migration rates using 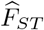. Contour plot shows the ability to rank between location pairs by 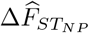 (horizontal axis), the absolute difference between 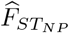 values for the two location pairs under comparison, and the *n*, the number of individuals sampled from each location (vertical axis). Each point represents the mean value for 100 independent values for PRC_*np*_, as described in the methods. *M* is a scaling factor for the distribution of pairwise migration rates between populations. *N*_e_ is the effective population size specified for each coalescent simulation.

